# From Channel-Pair Connectivity to Brain Networks: An Open Graph-Theoretical Pipeline for fNIRS Hyperscanning

**DOI:** 10.64898/2026.08.25.746918

**Authors:** Yael Hodaya Moshe, Mini Sharma, Anat Dahan, Hila Gvirts

## Abstract

Despite the growing use of functional near-infrared spectroscopy (fNIRS) hyperscanning to record brain activity simultaneously from interacting individuals in naturalistic settings, most analyses quantify functional connectivity separately for each channel pair. The resulting collection of pairwise estimates is difficult to integrate into a network-level characterization of intra- and inter-brain organization. Here, we present an open, configuration-driven Python toolkit that transforms preprocessed fNIRS hyperscanning time series into functional connectivity graphs. The toolkit constructs a bipartite inter-brain network for each dyad and separate intra-brain networks for each participant, computes node- and graph-level measures, and exports adjacency matrices, edge lists, analysis-ready summary tables, reproducibility metadata, and standardized visualizations. Dataset-specific parameters, including directory structure, participant naming, channel selection, epoch extraction, and edge-retention criteria, are defined in a human-readable YAML configuration file, enabling the same workflow to accommodate differently organized datasets without changes to the source code. We illustrate the pipeline using a representative recording from a mother-infant fNIRS hyperscanning dataset and present the resulting network outputs. The toolkit provides a reproducible framework for moving from pairwise functional connectivity estimates to network-level analyses of dyadic and individual brain organization.

## 1. Introduction

In recent years, social neuroscience has shifted from studying isolated brains toward examining neural activity as it unfolds during real, dyadic social interaction. Hyperscanning, the simultaneous recording of brain activity from two or more individuals, enables researchers to quantify inter-brain synchronization (IBS), which has been proposed to reflect mutual understanding and shared intentionality between interacting partners^1–5^.

Functional near-infrared spectroscopy (fNIRS) is particularly well suited to hyperscanning because it is portable, tolerant of naturalistic movement, and can be used with populations that are difficult to scan with fMRI, such as infants and young children^6–8^. The dominant analytic approach in fNIRS hyperscanning is wavelet transform coherence (WTC), which estimates the cross-correlation of two time series as a function of time and frequency, and can additionally be used to classify the directionality of coupling (in-phase, lagged, or anti-phase synchronization)^9^.

However, WTC and related coherence measures share a structural limitation: they characterize connectivity between a single pair of channels (or a single pair of regions) at a time, and do not by themselves describe how connectivity is organized across the whole set of recorded channels as a network. In single-brain neuroimaging, graph theory has provided a rich vocabulary (density, efficiency, centrality, modularity, and related metrics) for describing exactly this kind of network-level organization^10,11^. Extending this vocabulary to hyperscanning data requires representing each recording channel as a graph node and each statistically reliable association between channels as an edge. These nodes may belong to a single participant or to both members of a dyad.

Graph-theoretical analysis has become an increasingly important framework for characterizing large-scale brain organization from neuroimaging data, including EEG, MEG, fMRI, and fNIRS^12^. Rather than focusing on individual connections, graph representations capture the topology of the entire network, enabling the quantification of properties such as integration, segregation, efficiency, and hub organization^10,11^. More recently, these concepts have been extended to hyperscanning studies, where graph-based representations have been used to characterize both intra-brain and inter-brain functional networks and to investigate how network topology changes across different social and cognitive contexts. While these studies demonstrate the potential of graph-theoretical approaches for hyperscanning, most focus on specific datasets or custom analysis workflows rather than providing a general, reusable framework for constructing such networks from raw recordings^13^.

Despite the conceptual appeal of this approach, there is, to our knowledge, no widely available, open, reproducible pipeline that constructs both inter-brain and intra-brain connectivity graphs from raw hyperscanning recordings and outputs standard graph-theoretical metrics ready for group-level statistical analysis. Existing hyperscanning toolboxes largely focus on coherence/synchronization indices rather than graph construction^3,8,14^. Here, we present a configuration-driven, three-stage framework implemented as an installable Python package with a command-line interface. The pipeline takes participant-level hyperscanning recordings organized by dyad and session and performs three sequential stages: inspection, epoch extraction, and graph construction. The inspection stage reports available channels and flags missing files, duplicate or unrecognized participant identifiers, and missing channel columns. The extraction stage identifies task-relevant epochs according to configurable rules. The graph-construction stage generates inter-brain graphs for each dyad and intra-brain graphs for each participant, together with node- and graph-level metrics, adjacency matrices, edge tables, summary tables, and standardized visualizations. Because the pipeline operates purely on configured channel columns and a configured dyad/session folder structure, it is agnostic to the specific relationship between the two members of a dyad. The workflow is demonstrated here using a mother-infant fNIRS hyperscanning dataset.

## 2. Materials and Methods

An overview of the complete workflow is shown in Figure 1. The pipeline is implemented as a configuration-driven Python package, hyperscanning_toolkit (pandas, NumPy, SciPy, NetworkX^15^, PyYAML, Matplotlib), which is installable via pip and provides a command-line interface. It consists of three sequential stages - inspect, extract, and graphs, each of which reads the output of the previous stage and writes intermediate results to disk, so that the full pipeline is reproducible and each stage can be re-run or inspected independently. Every dataset-specific detail (folder layout, channel-column selection, epoch-extraction rule, and significance thresholds) is specified in a single YAML configuration file. Consequently, adapting the pipeline to a new dataset requires writing a new config file, not modifying source code.

**Figure 1.**
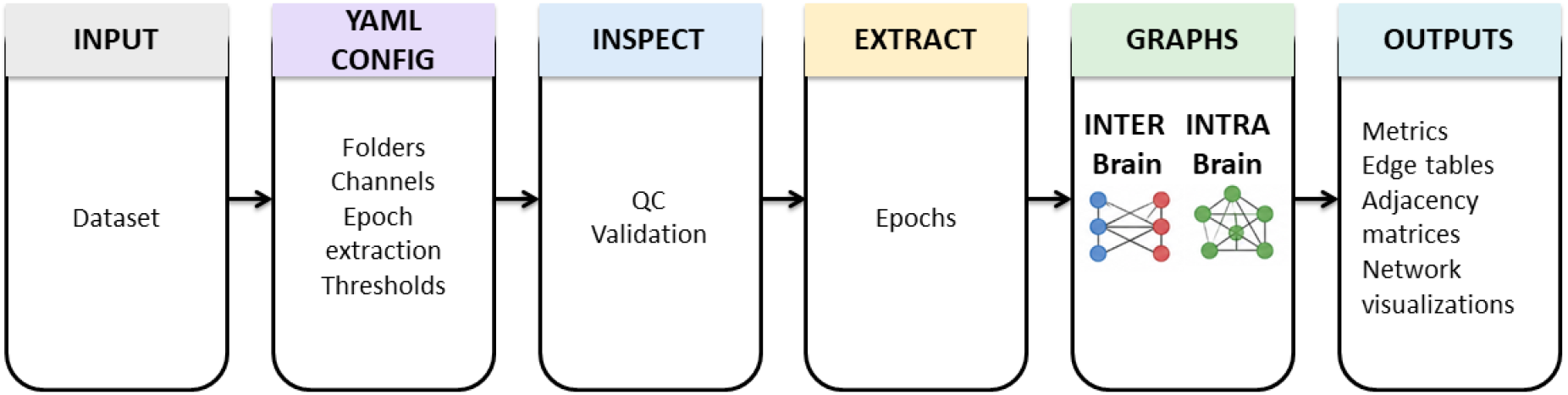
Workflow of the proposed hyperscanning toolkit. Dataset-specific parameters are provided through a YAML configuration file, enabling the same analysis pipeline to be applied to different hyper-scanning datasets without modifying the source code. The workflow includes dataset inspection, epoch extraction, graph construction, and automated export of network analysis results.

### 2.1 Data organization and channel inspection (inspect stage)

Directory structure, file naming, and channel-column selection are fully configurable through a YAML configuration file. A discovery rule identifies recording-session folders, while configurable filename patterns and regular expressions locate participant-specific files and extract subject identifiers. Channel columns can be selected automatically, by regular expression, or from an explicit column list. During the inspection stage, the toolkit scans the configured dataset, reports the number of rows, columns, and selected channels for each participant file, generates a summary table, and flags structural inconsistencies such as duplicate or unrecognized subject identifiers or missing channel columns. In the example dataset used to illustrate the toolkit, recordings are organized by dyad and experimental condition, with participant-specific files containing HbO channel measurements. The pipeline was also successfully applied to the complete example dataset, generating inter-brain and participant-specific intra-brain graphs for all available recordings.

### 2.2 Epoch extraction (extract stage)

Continuous recordings are restricted to the rows of interest for connectivity analysis using one of three configurable epoching modes: no filtering, in which the complete recording is retained; fixed-length windowing, in which recordings are divided into windows of a user-defined size; or event-marker epoching, in which data are extracted according to configurable start and end marker definitions. The epoching parameters, including window length or marker definitions, are specified once in the YAML configuration file and applied consistently to both participants in each dyad. This produces temporally aligned participant-specific epochs corresponding to the same experimental interval. The aligned epochs are subsequently used to calculate within-participant and cross-participant connectivity. The resulting connectivity matrices are used to construct the inter-brain and intra-brain networks described in Sections 2.3 and 2.4.

### 2.3 Inter-brain graph construction

For each dyad and extracted epoch, inter-brain connectivity is calculated by computing Pearson correlation coefficients between every channel from one participant and every channel from the other participant. Correlations satisfying user-defined significance and correlation thresholds (default: p < 0.05 and r > 0) are retained as edges. These thresholds are specified in the YAML configuration file. The resulting edge list is used to construct a weighted, undirected bipartite graph using the NetworkX library. Node-level metrics, including degree, node strength (weighted degree), betweenness centrality, closeness centrality, eigenvector centrality, and local clustering coefficient, are computed for every node. The toolkit computes both node-level and graph-level measures of inter-brain network organization. Node-level measures characterize the contribution of individual channels to cross-participant connectivity, whereas graph-level measures characterize the overall organization of the dyadic network. Graph-type-specific measures, including bipartite density, account for the division of nodes between the two 2.4 Intra-brain connectivity graph construction (graphs stage - intra-brain).

For each participant and extracted epoch, intra-brain connectivity is calculated independently among the channels belonging to that participant. Pearson correlation coefficients are computed between all unique pairs of channels within a participant (i.e., each pair of channels is considered only once), and channel pairs satisfying user-defined significance and correlation thresholds are retained as graph edges. This results in one weighted, undirected intra-brain graph for each participant and. The toolkit computes the same core set of node-level and graph-level measures for both intra-brain and inter-brain networks. However, these measures have different interpretations across the two graph types: in intra-brain graphs, they characterize connectivity among channels within an individual participant’s brain, whereas in inter-brain graphs, they characterize connectivity between channels belonging to different participants. Intra-brain analyses additionally include modularity, which quantifies the division of the network into communities, whereas inter-brain analyses include graph-type-specific measures such as bipartite density.

### 2.4 Intra-brain connectivity graph construction

For each participant and extracted epoch, intra-brain connectivity is calculated independently among the channels belonging to that participant. Pearson correlation coefficients are computed between all unique pairs of channels within a participant (i.e., each pair of channels is considered only once), and channel pairs satisfying user-defined significance and correlation thresholds are retained as graph edges. This results in one weighted, undirected intra-brain graph for each participant and extracted epoch. The toolkit computes a shared core set of node- and graph-level measures for both network types. However, these measures have different interpretations across the two graph types: in intra-brain graphs, they characterize connectivity among channels within an individual participant’s brain, whereas in inter-brain graphs, they characterize connectivity between channels belonging to different participants. Intra-brain analyses additionally include modularity, which quantifies the division of the network into communities, whereas inter-brain analyses include graph-type-specific measures such as bipartite density. The complete list of shared and graph-type-specific metrics and their definitions is provided in Table 1.

**Table 1.** Graph-theoretical metrics exported by the Hyperscanning Toolkit. Node-level metrics characterize individual recording channels, whereas graph-level metrics summarize the overall topology of the connectivity network. The Graph Type column indicates whether each metric is computed for inter-brain graphs, intra-brain graphs, or both. *† Note*. *Standard triangle-based clustering measures are not informative for a pure bipartite graph*.

| Metric | Graph Type | Description |
| --- | --- | --- |
| **Node-level metrics** |  |  |
| Degree | Both | Number of retained edges connected to a node, indicating how many other channels are connected to that channel. |
| Node strength (weighted degree) | Both | Sum of the weights of all retained edges connected to a node, reflecting the overall strength of its connections. |
| Degree centrality | Both | Degree normalized by the maximum possible number of connections for the relevant graph type. |
| Betweenness centrality | Both | Measures how frequently a node lies on the shortest paths between other nodes, indicating its potential role as a bridge within the network. |
| Closeness centrality | Both | Measures the proximity of a node to all other reachable nodes based on shortest-path distances. Higher values indicate greater network accessibility. |
| Eigenvector centrality | Both | Measures node centrality by considering both its connections and the centrality of the nodes to which it is connected. |
| Local clustering coefficient | Intra-brain only† | Measures the extent to which the neighboring nodes of a given node are also interconnected. |
| **Graph-level metrics** |  |  |
| Number of nodes | Both | Total number of nodes, representing recording channels, included in the graph. |
| Number of edges | Both | Total number of retained connections in the graph. |
| Density | Intra-brain only | Fraction of observed within-participant edges relative to all possible within-participant edges. |
| Mean positive edge weight | Both | Mean correlation strength across all retained positive edges. |
| Maximum edge weight | Both | Largest edge weight in the graph. |
| Minimum positive edge weight | Both | Smallest positive edge weight among the retained edges. |
| Average node strength | Both | Mean weighted degree across all nodes in the graph. |
| Average degree | Both | Mean number of retained connections per node. |
| Average clustering coefficient | Intra-brain only† | Mean local clustering coefficient across all nodes, reflecting the overall tendency to form local clusters. |
| Global efficiency | Both | Quantifies network integration based on the inverse shortest-path distances between nodes. |
| Number of connected components | Both | Number of disconnected subnetworks within the graph. |
| Largest connected component size | Both | Number of nodes contained in the largest connected subnetwork. |
| Transitivity | Intra-brain only† | Global clustering measure based on the proportion of connected triples that form triangles. |
| Assortativity coefficient | Both | Measures whether nodes tend to connect to other nodes with a similar degree. |
| Bipartite density | Inter-brain only | Fraction of observed cross-participant edges relative to all possible cross-participant edges. |
| Number of connected group A nodes | Inter-brain only | Number of nodes from participant A with at least one retained inter-brain connection. |
| Number of connected group B nodes | Inter-brain only | Number of nodes from participant B with at least one retained inter-brain connection. |
| Modularity | Intra-brain only | Measures the extent to which the network can be partitioned into communities with relatively dense within-community and sparse between-community connections. |

### 2.5 Software availability

The proposed pipeline is implemented in Python 3 as the installable package hyperscanning_toolkit (version 1.0.0). The package uses widely adopted scientific Python librarie, including pandas, NumPy, SciPy, NetworkX, PyYAML, and Matplotlib, and provides a command-line interface supporting the inspect, extract, graphs, and run commands. To ensure reproducibility, each analysis automatically records execution metadata, including the toolkit version, Git commit hash, execution timestamp, configuration file, analysis parameters, and software environment in a run_metadata.json file generated alongside the analysis outputs. The package also includes automated tests to verify correct functionality across configurable dataset organizations and analysis settings. The source code is publicly available under the MIT License at: https://github.com/yaelhm/hyperscanning-toolkit

## 3. Illustrative Application and Outputs

To demonstrate the capabilities of the toolkit, we analyzed preprocessed HbO recordings from a representative mother-infant dyad, identified in the dataset as dyad 100, during the Elicit (E) condition, in which the mother and infant performed a semi-structured cooperative task designed to naturally evoke interpersonal synchrony through shared goals (e.g., joint tower building). The recordings of both participants contained 1,892 aligned time points, which served as input for connectivity estimation, graph construction, and network analysis.

**Figure 2.**
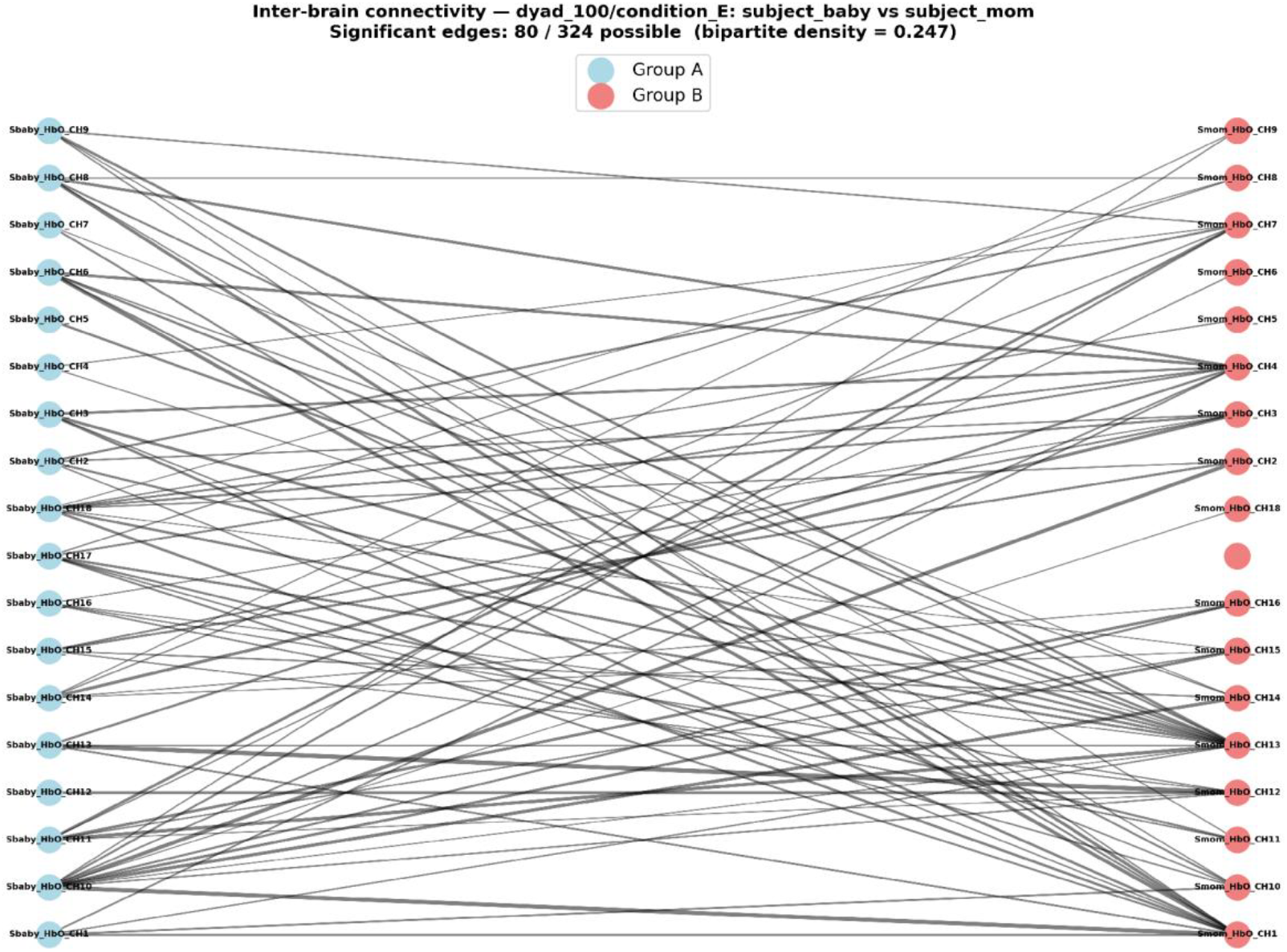
Inter-brain (bipartite) connectivity graph, for dyad 100 during the Elicit condition (infant-mother), computed from 18 HbO channels per participant. Node color denotes participant (blue = infant, red =mother). Edges represent cross-participant channel pairs that met the configured Pearson correlation thresholds (p < 0.05, r > 0) Pearson correlation across the two brains, with width proportional to r. In this example, 80 of 324 possible infant - mother channel pairs were retained corresponding to a bipartite density of 0.247).

**Figure 3.**
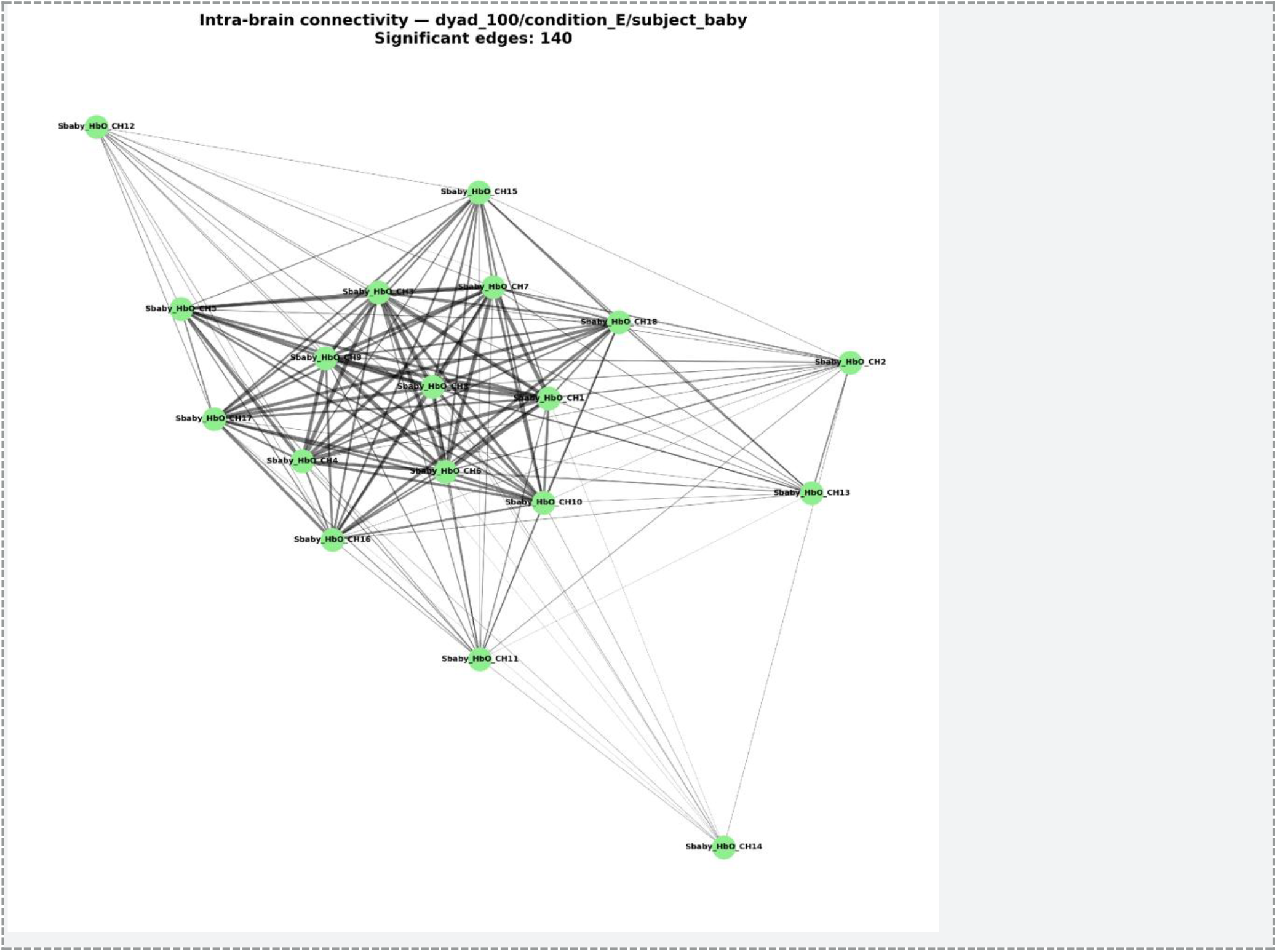
Infant intra-brain connectivity graph for dyad 100 during the Elicit condition, computed from 18 HbO channels. Edges represent within-participant channel pairs that met the configured Pearson correlation thresholds (p < 0.05 and r > 0), with edge width proportional to r. In this example, 140 of 153 possible within-participant channel pairs were retained, corresponding to a density of 0.915.

**Figure 4.**
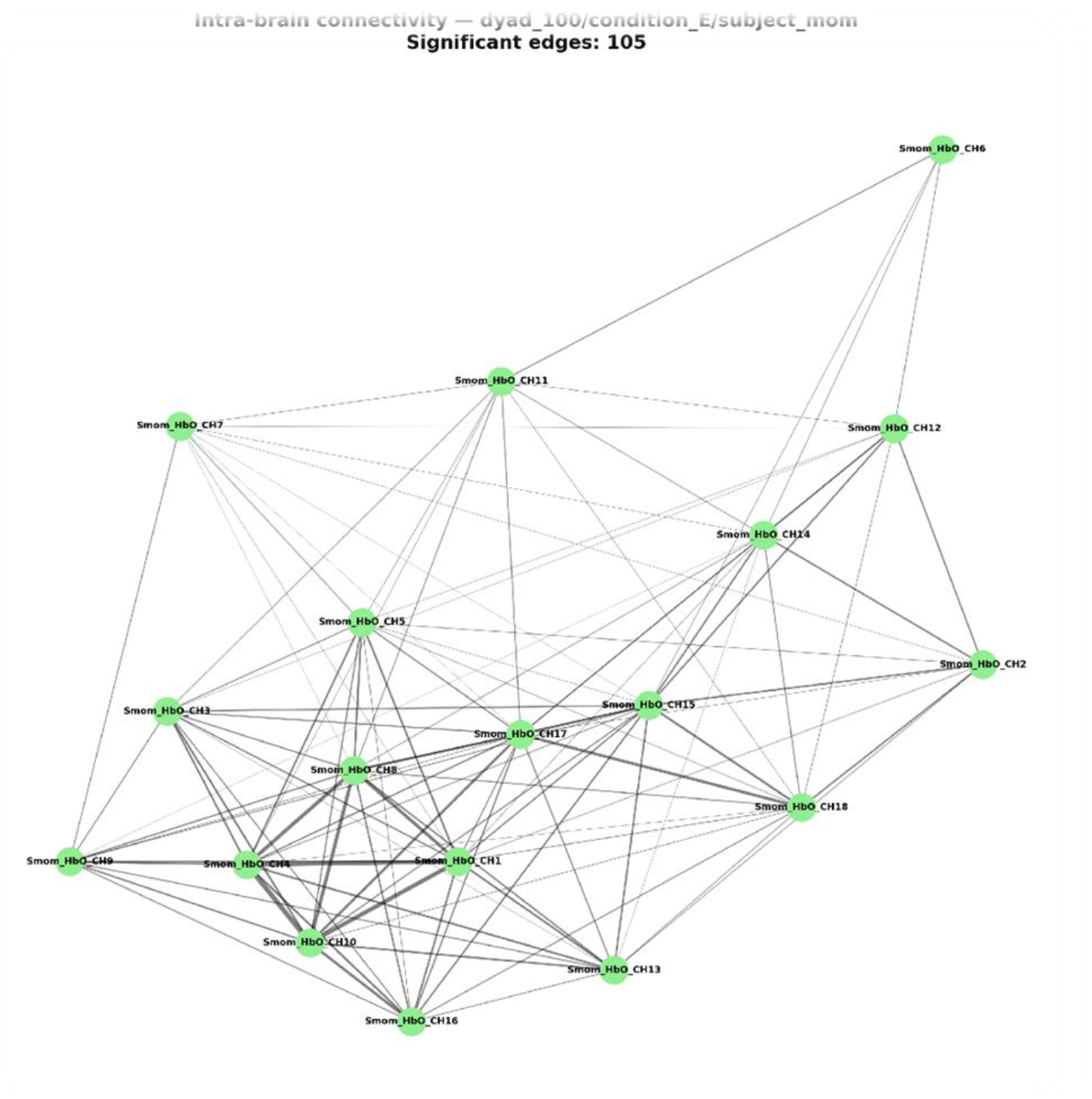
Maternal intra-brain connectivity graph for dyad 100 during the Elicit condition, computed from 18 HbO channels. The same visualization conventions and edge-retention criteria described for Figure 3 were applied. In this example, 105 of the 153 possible within-participant channel pairs were retained, corresponding to a density of 0.686.

For the illustrative inter-brain network (Figure 2; Table 2), 80 of the 324 possible infant-mother channel pairs were retained after thresholding, resulting in a graph with a bipartite density = 0.247. The network had average node strength of 0.358, an average degree of 4.44, and a global efficiency of 0.447. It consisted of two connected components, with the largest containing 35 of the 36 nodes, and exhibited negative degree assortativity (-0.260). All 18 infant channels and 17 of the 18 maternal channels participated in at least one significant inter-brain connection. In this illustrative network, the two nodes with the highest strength values were maternal channels, with Smom_HbO_CH13 (node strength = 1.197) and Smom_HbO_CH1 (1.079) showing the highest strengths, followed by Sbaby_HbO_CH10 (1.000).

**Table 2.** Summary of graph-level metrics for the illustrative inter-brain and intra-brain networks shown in Figures 2-4.

| Metric | Value |
| --- | --- |
| Inter-brain (infant×mother) |  |
| Nodes / retained edges (of possible) | 36 / 80 (of 324) |
| Bipartite density | 0.247 |
| Global efficiency | 0.447 |
| Average node strength | 0.358 |
| Average degree | 4.44 |
| Intra-brain (infant) |  |
| Nodes / retained edges (of possible) | 18 / 140 (of 153) |
| Density | 0.915 |
| Average clustering coefficient | 0.386 |
| Global efficiency | 0.958 |
| Modularity | 0.028 |
| Intra-brain (mother) |  |
| Nodes / retained edges (of possible) | 18 / 105 (of 153) |
| Density | 0.686 |
| Average clustering coefficient | 0.203 |
| Global efficiency | 0.843 |
| Modularity | 0.176 |

The intra-brain networks (Figures 3-4; Table 2) illustrate the participant-specific outputs generated by the toolkit. In this example, the infant network was denser than the maternal network (density = 0.915 vs. 0.686), and had a higher average clustering coefficient (0.386 vs. 0.203), global efficiency (0.958 vs. 0.843), and average node strength (6.09 vs. 2.56). Modularity was lower in the infant network (0.028) than in the maternal network (0.176), descriptively reflecting the nearly complete structure of the infant graph and the comparatively greater modular organization of the maternal graph in this illustrative recording. The strongest nodes were Sbaby_HbO_CH8 (node strength = 9.59) for the infant and Smom_HbO_CH10 (node strength = 4.42) for the mother.

## 4. Discussion

The pipeline extends conventional pairwise hyperscanning analyses by transforming channel-level Pearson-correlation estimates into graph-based representations of network organization. It constructs two complementary types of networks: a dyadic inter-brain network, in which edges represent connectivity between channels belonging to different participants, and participant-specific intra-brain networks, in which edges represent connectivity among channels within each participant’s brain. By combining shared metrics with graph-specific measures, the toolkit supports complementary descriptions of dyadic and participant-level network organization.

A central design principle of the toolkit is flexibility across hyperscanning datasets. Dataset-specific parameters, including directory structure, file naming conventions, channel selection, epoch extraction, and statistical thresholds are defined in a single YAML configuration file, allowing the same analysis pipeline to be applied to datasets with different organizational structures without modifying the source code. In this study, the complete workflow is demonstrated using a mother-infant fNIRS hyperscanning dataset.

Consequently, the toolkit is suitable for a wide range of hyperscanning paradigms, including mother-infant, caregiver-child, therapist-client, and other dyadic interaction studies that produce simultaneously recorded fNIRS time series from two participants.

## 5. Limitations and Future Directions

The current implementation has several limitations. First, connectivity is estimated using static Pearson correlation across the complete recording epoch and therefore does not capture temporal changes in network organization. Subsequent versions will incorporate sliding or adaptive time windows to support time-varying connectivity analysis. Second, edges are retained using configurable significance and correlation thresholds, and the influence of multiple-comparison correction requires systematic evaluation. Alternative thresholding strategies will therefore be examined. Finally, Pearson correlation quantifies statistical association but does not estimate directed information flow. Additional connectivity measures, including wavelet transform coherence and other functional-connectivity indices, are planned for future releases.

The present illustration used a representative recording from a mother–infant fNIRS hyperscanning dataset. Further evaluation across additional datasets and experimental paradigms will help establish the robustness and generalizability of the toolkit.

